# Targeting the G-quadruplex structure in the hTERT promoter: In silico screening of phytocompounds and replica exchange molecular dynamics simulations

**DOI:** 10.1101/2023.12.31.573762

**Authors:** Akshay Uttarkar, Vidya Niranjan

## Abstract

Telomerase activity plays a crucial role in maintaining telomere length and cellular immortality, making it an attractive target for cancer therapy. The human telomerase reverse transcriptase (hTERT) promoter contains a G-rich region that can form G-quadruplex (G4) structures, which have been shown to regulate hTERT expression. In this study, we used in silico screening and molecular dynamics simulations to identify phytocompounds that can stabilize the G4 structure in the hTERT promoter. We performed shape-based and pharmacophore-based screening of a phytochemical database and identified two lead compounds with assistance from oleanolic acid and maslinic acid as controls which showed in vitro telomerase activity. Molecular docking and replica exchange molecular dynamics simulations for a temperature profile of 300K to 350K were used to evaluate the binding affinity and stability of these compounds with two different G4 conformations in the hTERT promoter. Our results suggest that astragaloside-1 can stabilize the parallel-stranded G4 conformation (2kze) in the hTERT promoter, while novel compounds may be required to stabilize the intramolecular G4 conformation (2kzd). Our study highlights the potential of in silico screening and molecular dynamics simulations in identifying lead compounds for targeting G4 structures.

## 1. Introduction

Telomerase, a ribonucleoprotein complex, is crucial for maintaining telomeres, the protective structures at the ends of chromosomes. It is known to add telomeric repeats during the replicative phase of the cell cycle[1]. Research has shown that telomerase activity is closely linked to the cell cycle, with the highest levels detected in S-phase cells, while cells arrested at the G2/M phase exhibit significantly lower telomerase activity[1]. Various cell cycle blockers, such as transforming growth factor beta1 and cytotoxic agents, have been found to inhibit telomerase activity, establishing a direct association between telomerase activity and cell cycle progression[2]. Additionally, induction of quiescence by serum deprivation does not impact telomerase activity, and cells arrested at the G1/S phase show similar levels of telomerase activity to those in the S phase[3]. This modulation of telomerase activity throughout the cell cycle highlights its dynamic regulation and its importance in cell proliferation and cancer development. The specific association of telomerase activity with immortal cells and cancer has also been documented in previous studies[4,5]. These findings underscore the significance of understanding telomerase activity in the context of the cell cycle and its potential implications for cancer research and therapy.

Telomerase activity is regulated by various factors, including the cell cycle, genetics, and environmental stressors. Telomerase activity is closely linked to the cell cycle, with the highest levels detected in S-phase cells, while cells arrested at the G2/M phase exhibit significantly lower telomerase activity[1]. Various cell cycle blockers, such as transforming growth factor beta1 and cytotoxic agents, have been found to inhibit telomerase activity, establishing a direct association between telomerase activity and cell cycle progression[6]. Genetic mutations or polymorphisms in TERC and/or TERT can increase telomerase activity, leading to telomere lengthening[2]. Additionally, oxidative stress and inflammation can cause DNA damage and telomere shortening, which can be replenished by telomerase[2,4]. Other factors that can affect telomerase activity include demographics, such as age and sex, and metabolic changes[3,4]. Telomerase activity is mainly present in stem, sex, and cancer cells, while most somatic cells lack or have very minimal telomerase activity[7]. The stimulation of telomerase activity can prevent or delay cellular aging triggered by critical telomere shortening[4,5]. These findings highlight the complex regulation of telomerase activity and its potential implications for aging and disease.

Several factors can affect telomerase activity, with human telomerase reverse transcriptase (hTERT) playing a central role. hTERT expression is tightly regulated at the transcriptional level and is closely associated with telomerase activity, making it the primary determinant for the enzyme activity[8]. The regulation of hTERT expression involves various mechanisms, including pre-mRNA alternative splicing, genetic and epigenetic alterations, and hormonal control[8–10]. Additionally, several transcription factors, such as c-Myc, Max, Sp1, and Ets1, are implicated in hTERT expression, acting as activators in some cases[11]. Furthermore, telomerase activity and hTERT expression are under the control of steroid sex hormones and growth factors in hormonally regulated tissues and certain cancers[12]. The complex regulation of hTERT and telomerase activity underscores the multifaceted nature of their control, involving genetic, epigenetic, and hormonal factors. Understanding these regulatory mechanisms is crucial for elucidating the role of telomerase in cancer and aging and for the development of potential therapeutic interventions.

G-quadruplexes (G4s) are unique noncanonical DNAs that play a key role in cellular processes and neoplastic transformation[13]. The hTERT promoter, which contains many G-tracts on the same DNA strand, exhibits an exceptional potential for G-quadruplex formation[14,15]. The formation of G-quadruplexes in the hTERT promoter has been shown to inhibit hTERT expression, leading to telomere shortening and senescence[16]. Recent studies have also suggested that G-quadruplexes in the hTERT promoter can play a role in hTERT activation[13–15]. The coexistence of two distinct G-quadruplex conformations in the hTERT promoter has been reported, with one of them being more stable and potentially involved in hTERT activation[13]. Additionally, mutations in G-tracts in the hTERT promoter have been shown to abrogate G4-folding and activate hTERT expression, suggesting a potential target for cancer therapy[17]. The regulation of hTERT expression and telomerase activity by G-quadruplexes in the hTERT promoter underscores the importance of understanding the complex regulatory mechanisms of telomerase in cancer and aging.

Small molecules that promote G-quadruplex formation in the hTERT promoter can lead to the upregulation of hTERT expression in cancer cells. However, the majority of research has focused on the inhibition of hTERT expression by stabilizing the higher-order hTERT G-quadruplex with small molecules[18–21]. For instance, a disubstituted 2-aminoethyl-quinazoline has been shown to selectively bind across the G4 junctional sites of the hTERT promoter, resulting in the downregulation of hTERT transcription in breast cancer cells[18]. Similarly, compounds like TMPyP4, Telomestatin, and GTC365 have been shown to stabilize the hTERT G-quadruplex, offering potential for the indirect inhibition of telomerase and the development of novel anticancer therapeutics[18, 21]. However, recent studies have suggested that small molecules can also promote G-quadruplex formation in the hTERT promoter, leading to the upregulation of hTERT expression[18]. These small molecules have been investigated for their ability to target the G4 structure in the hTERT promoter, providing evidence that modulating the G4 structure with small molecules is a viable approach to modulate telomerase activity and hTERT expression[22]. Further research is needed to elucidate the role of small molecules in promoting G-quadruplex formation in the hTERT promoter and their potential as therapeutic targets.

The current focus of the research work is to screen the phytocompounds for its ability to stabilize higher order of G-quadruplex complexes via Replica exchange molecular dynamics (REMD) simulation leading to activation and expression of hTERT and compare its activity with available control compounds which have shown invitro telomerase activity.

## 2. Methods

### 2.1. Selection of G-quadruplex DNA structure and phytocompounds

ONQUADRO [23] is a comprehensive database providing experimental structures of G-quadruplex. For hTERT proteins two structures were available reposited in RCSB-PDB [24] bearing the id’s 2KZE [25] and 2KZD [26]. 2KZD (Fig 1A) is a structure of a (3+1) G-quadruplex formed by hTERT promoter sequence whereas 2KZE (Fig 1B) is a structure of an all-parallel-stranded G-quadruplex formed by hTERT promoter sequence. Both the structures have 10 conformers submitted. These two structures were selected for molecular docking studies.

**Figure 1.**
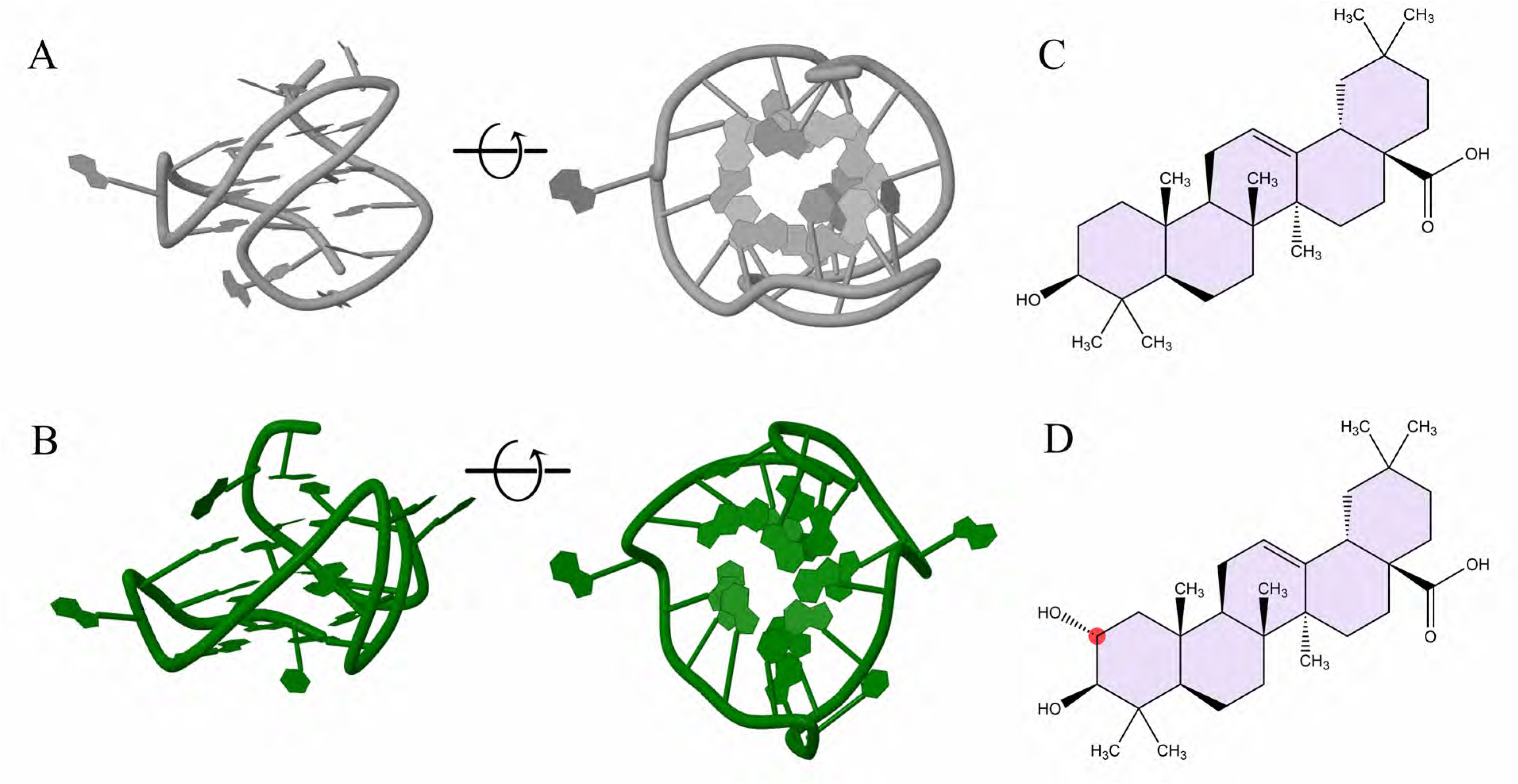
Structural Information of Quadruplex Structures and query compounds. A) DNA G-4 quadruplex structures 2KZD conformation showing in the minor and major grooves which is rotated on x-axis to show the guanine rich double helix structures. B) DNA G-4 quadruplex structures 2KZE conformation showing in the minor and major grooves which is rotated on x-axis to show the guanine rich double helix structures. C) 2D fischer structure of Oleanolic acid is terminal carbons shown and rings coloured in grey. D) 2D fischer structure of Maslinic acid is terminal carbons shown and rings coloured in grey. The C27 is shown in Red.

The phytocompounds used in the study are an amalgamation of compounds from MolPort, Coconut [27] and Super-Natural [28]. A total of 380,000 compounds are used to create an in-house database of phytocompounds. These include both Natural and Natural Like compounds.

The compounds were subjected to ligand preparation [29] via minimization for 500 steps and generate 32 conformers for each compound. The best conformer for each compound is selected for screening.

For the current study, we have selected two phytocompounds which have shown invitro telomerase activity which are Oleanolic acid [30] (Fig 1C) and Maslinic acid [31] (Fig 1D). These structures are downloaded from PubChem [32].

### 2.2. 1-D based and pharmacophore-based screening of Phyto database

First, a shape data file needs to be created for 79710 compounds providing 3,815,713 conformers. A shape data creation module [33] in Schrodinger was used. The uniqueness of each ligand was set to stereoisomers and out conformer set to 1. The shape data was calculated and stored as a typed pharmacophore which would assist in the next steps of analysis.

The molecules were screened based on the control compounds. In can we be observed that, maslinic acid has an additional -OH group via single down bond at position C27 compared to oleanolic acid. This makes the cleared that the size and volume along with the reactive groups in both the compounds is significantly similar.

With control compounds as a template the phytocompounds shape data created was screened with similarity set to >0.90 and similarity normalization set to query compounds.

The control compounds (query) were used to build a pharmacophore hypothesis [34]. Both the compounds were set to be active. The features under consideration were Acceptor, Donor, Hydrophobic sites, Negative and Positive Ionic sites and Aromatic rings. The model is developed based on alignment and common features. The best pharmacophore model was used to screen the compounds obtained from 1-D screening.

### 2.3. Structural fingerprint-based clustering of compounds

Atom fingerprinting is a quick and effective method to cluster the compounds based on the various atom features. In the current study, canvas similarity and clustering module of Schrodinger was used to perform the analysis. The precision was set to 64 bit and fingerprint type set to radial with atoms distinguished based on terminal, halogen, hydrogen bond acceptor/donor, and its bond order along with hybridization.

Tanimoto clustering method was used alpha and beta values set to 0.5.

The linkage method was set to average and the best cluster was used for virtual docking.

### 2.4. Molecular docking of lead compounds with DNA G-quadruplex

#### 2.4.1. Quadruplex docking preparation

The PDB structures 2KZD and 2KZE was imported into Maestro [35] window. The protein preparation module was used to check for missing atoms and minimizing the structure. OPLS3 [36] force field is added to the quadruplex system. Sitemap [37,38] is used to identify potential sites in the DNA groves for docking of compounds to stabilize the structure. Using the site available for docking a receptor grid was developed for docking with default values.

#### 2.4.2. Extra precision based molecular docking

The final compound from the cluster is subjected to extra precision docking [39] using Glide [40,41]. Along with the phytocompounds the query compounds were also subjected to molecular docking for comparison. The scaling factor and partial cutoff charge was set to 0.80 and 0.15 respectively. The ligand sampling was set to flexible. No excluded volume penalty was applied, and no constraints were added for molecular docking.

### 2.5. Replica Exchange based MD simulation

REMD simulations were running using Desmond [42], Maestro. System Builder under Desmond was used to create the simulation system and the solvent model used was Transferable intermolecular potential 3P (TIP3P) [43,44]. An orthorhombic box shape was used to define the boundary conditions and the volume was minimized to enclose the complex. The system was not neutralized by the addition of Na+ or Cl-. The force field applied was OPLS3. MD simulations were carried out for a simulation time of 100ns with a recording interval of 0.1ns.

The tempering method used was Parallel REMD for a temperature of 300K to 350K. The specific range was selected based on the melting temperature calculated using Oligo calc [45]. A linear temperature profile was followed with temperatures at 300K, 312K, 325K, 337K, and 350K. The simulation time was 100 nanoseconds with NVT as the set ensemble class. The other parameters were same as that of MD simulations mentioned the previous section 2.3.

The principal component analysis (PCA) was carried out using Evol [46] and Prody [47] tools using VMD [48].

## 3. Results

### 3.1. Shape based 1-D screening for improved efficient screening with query compounds

1D-fingerprint comparison to quickly assess the similarity between the reference and the library molecules without the need for full 3D molecular alignment, and rapidly generates a similarity score for each molecule.

The algorithm treats all atoms equivalently and computes the overlap between structures A and B as the sum of pairwise atomic overlaps, normalized by the largest self-overlap.

The compounds from various phytocompounds databases were downloaded as mentioned in the methods section. The duplicates from were verified based on smiles-based screening and the unique compounds were used for ligand preparation.

A total of 79710 compounds were available for screening which were subjected to 1D screening after completion of 1D data bin database.

For oleanolic acid, as a query compound the aligned structures were 159,422 and similarly for maslinic acid it was 318,846. It can be strongly seen that even though both these compounds differ only by a -OH group, they alignment conformers are increased by 200% for maslinic acid.

This highlights the robustness of the screening parameters set and its ability to screen and shortlist compounds from a large database. Overall concatenated conformers for screening are 478,266 compounds.

### 3.2. Pharmacophore based screening funnels the hit phytocompounds

Generation of pharmacophore hypothesis/model creates a 3D geometry-based screening of by aligning both Oleanolic Acid and Maslinic acid. Essential spatial and electronic characteristics necessary for a molecule is generated. These essential features are hydrogen bond acceptors (HBA), hydrogen bond donors (HBD), aromatic rings, and hydrophobic groups.

A total of 20 pharmacophores models were generated. Each model was classified and ranked based on the model accuracy score. A merged hypothesis was generated to obtain a consensus of all the generated models. In this final model generated the features of importance were 1 Acceptor, 1 Donor and 3 Hydrophobic sites.

This model was used to screen the database of 478,266 conformers of which 15,492 had a perfect 5 on 5 matches of features. A scatter plot with Fitness versus Align score of the compounds coloured on the basis of Volume mapped to pharmacophore is provided in Fig 2A.

**Figure 2:**
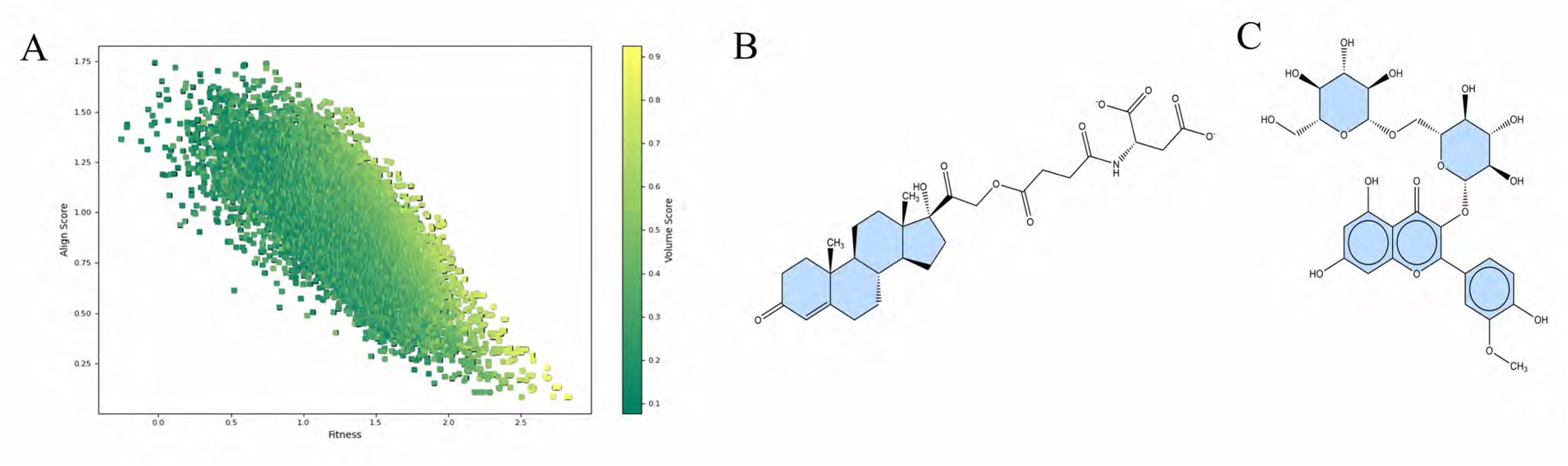
Screening of compounds and lead compounds from molecular screening. A) A 2D profile of compounds screened from pharmacophore with Fitness versus Align score with points coloured based on Volume score of the compounds. B) 2D fischer structure of preganone derivative is terminal carbons shown and rings coloured in sky blue. C) 2D fischer structure of Astragaloside-1 is terminal carbons shown and rings coloured in sky blue.

These shortlisted compounds were carried forward for further screening.

### 3.3. Clustering of compounds to obtain best lead phytocompounds

Clustering of compounds in drug screening is crucial for identifying similar molecules, reducing the computational burden, and enhancing hit confirmation rates. By grouping compounds based on their chemical properties, clustering enables the identification of active compounds and the exploration of chemical space, expediting the drug discovery process. Additionally, cluster-based enrichment strategies have been shown to improve hit confirmation rates, making them valuable in settings with limited confirmatory screening capacity. Furthermore, clustering allows for the analysis of compound diversity and the prioritization of structurally similar compounds, aiding in lead selection and drug design.

In the current work we cluster compounds based on their molecular properties, such as shape, electrostatics, and chemical features. The clustering process involves calculating the pairwise similarity between compounds and grouping them based on their similarity scores. Atoms were distinguished based on terminal, Hydrogen bond donors/acceptors, bond order and bond hybridization.

Off the 15,492 compounds, a total of 34 clusters were formed with a cluster strain of 1.00. These clusters were ranked based on the range of the volume score they acquire and fitness with the query compounds. Based on these filtering criteria, a cluster with 68 compounds was fit for further processing.

### 3.4. Molecular docking and identification of best compounds

The 68 compounds are now considered as hit molecules. These hit molecules were docked against two targets which are 2kzd and 2kze. The selection of these targets is previously mentioned in the methods section. These molecules were docked against DNA quadruplex targets. With 2kzd 98.5% compounds showed binding affinity and 97% with 2kze.

The best docked compound with 2kzd is a compound of IUPAC name (2S)-2-[[4-[2-[(8R,9S,10R,13S,14S,17R)-17-hydroxy-10,13-dimethyl-3-oxo-2,6,7,8,9,11,12,14,15,16-decahydro-1H-cyclopenta[a]phenanthren-17-yl]-2-oxoethoxy]-4-oxobutanoyl]amino]butanedioic acid (Fig 2B) bearing PubChem CID 67798210 [49] with a docking score of -8.78 kcal/mol. Currently there are no synonyms available for this compound but they belong to the super class of lipids, class of steroids and sub class of pregnane steroids. Its direct parent is glucocorticoid or progestogins. For the ease of labelling this compound henceforth will is labelled as pregnane-derivative. For the ease of naming convention in the manuscript, the docked complex with 2kzd, this will be henceforth mentioned as complex 1. The compound is docked in the major groove/bend in the (3+1) G-quadruplex. The interacting nucleotides are G19. The nucleotides in the biding groove are A6, G7, G8, G15, G18 and C20. The interacting pocket in the guanine strand.

Similarly, for 2kze it is Astragaloside-1 bearing PubChem ID 13996685 [50] (Fig 2C) with a docking score of -5.649 kcal/mol. Even these belong to the super class of lipids, class of steroids and sub class of Steroidal glycosides. Direct parent is cucurbitacin glycoside. This docked complex with 2kze will be referred to as complex 2. Astragaloside-1 is interacting with G5, G8, G9 and G15. The nucleotides part of the binding grove A6, G7 and G19. The interaction is more stable and forms a “bridge” kind of a structure to stabilize the structure. More details on the changes in conformation and stability can be obtained via simulation studies.

2D interaction profile of complex 1 is provided in Fig 3A and that of complex 2 is provided in Fig 3B.

**Figure 3:**
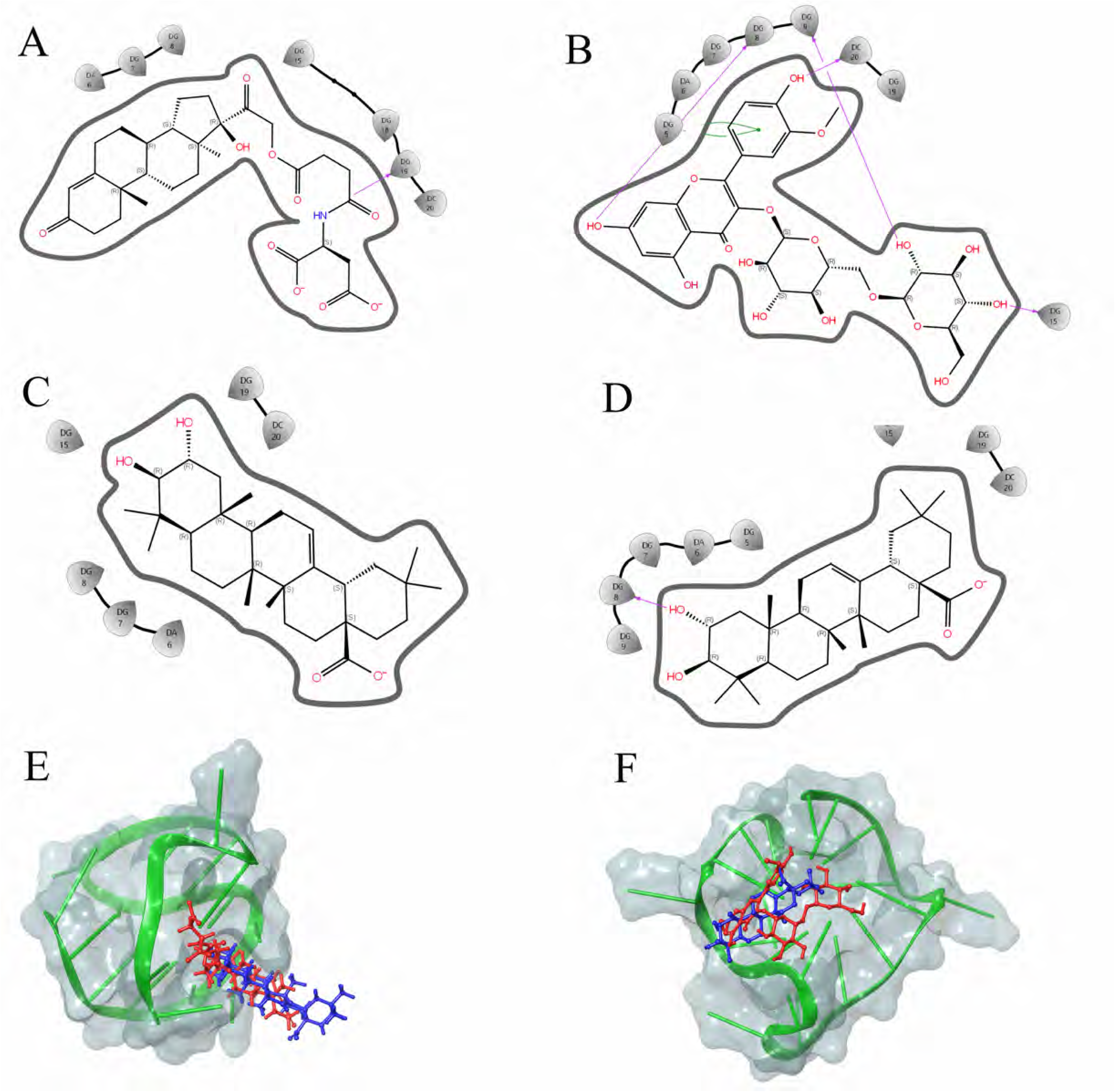
2D interaction profile and 3D binding pose of control and lead compounds. A) 2D interaction profile of preganone derivative with 2kzd nucleotides highlighted. B) 2D interaction profile of Astragaloside-1 with 2kze nucleotides highlighted. C) 2D interaction profile of Maslinic acid with 2kzd nucleotides highlighted. D) 2D interaction profile of Maslinic acid with 2kze nucleotides highlighted. E) A comparison of pregnane derivative (RED) with maslinic acid (BLUE) is provided. The DNA (GREEN) is shown with transparent surface visualization (Turquoise). F) A comparison of Astragaloside-1 (RED) with maslinic acid (BLUE) is provided. The DNA (GREEN) is shown with transparent surface visualization (Turquoise).

The query compounds were also subjected to molecular docking and maslinic acid was showed better binding efficiency with 2kzd and 2kze with a docking score of -6.23 kcal/mol and -6.56 kcal/mol respectively.

For the sake of easing out naming conventions, maslinic acid with 2kzd is referred to as control 1 and maslinic acid with 2kze as control 2.

2D interaction profile of control 1 is provided in Fig 3C and that of control 2 is provided in Fig 3D.

In Fig 3E, a comparison of pregnane derivative (RED) with maslinic acid (BLUE) is provided. The DNA (GREEN) is shown with transparent surface visualization (Turquoise). The control and compound superimpose each other in the binding groove. Similarly in Fig 3F, a comparison of Astragaloside-1 (RED) with maslinic acid (BLUE) is provided. The DNA (GREEN) is shown with transparent surface visualization (Turquoise). The control and compound superimpose each other in the binding groove.

These 4 complexes which are complex 1, complex 2, control 1 and control 2 along with two native DNA quadruplex structures are subjected to replica exchange based dynamic studies.

### 3.5. Replica exchange dynamics studies for understanding the quadruplex conformation in various conditions

REMD simulations have been employed to overcome high-energy barriers and efficiently sample the conformational space of biomolecular systems, making them valuable for understanding the dynamic behaviour of DNA quadruplex structures in the presence of drugs. These simulations enable the exploration of the free energy landscape of biomolecular systems, providing insights into the conformational ensembles and dynamic properties of DNA quadruplexes in complex with drug molecules [51].

In the current study, REMD was performed for all the 6 cases as mentioned in the previous section. The temperature range and steps along with the other parameters are provided in the methods section.

#### 3.5.1. REMD studies in native quadruplex

The DNA quadruplex structures considered for the study, 2kzd and 2kze are subjected to simulations to obtain insights on the folding and conformation pattern of the nucleotides. The percentage of accept ratio was 0.24 which is in the accepted range.

RMSD of the nucleotides was monitored throughout the simulation period.

A principal component analysis was performed to have a consensus in the change in structure and fluctuation occurring in the DNA quadruplex structure.

From Fig 4A, a structural fluctuation across the temperature profiles is shown as a box plot with a cut in the mean position. As well-known conventions, with rise in temperature the structural deviations increase and the same in provided in table 1. A cross correlation plot at each respective temperature profile from 300 to 350K is shown in Fig 4B to 4F respectively. From 325K onwards the structural profile looks similar and fairly overlapping is nature. The RMSD values are comparable and no statistically significant differences (P < 0.05) between various temperature profiles.

**Figure 4:**
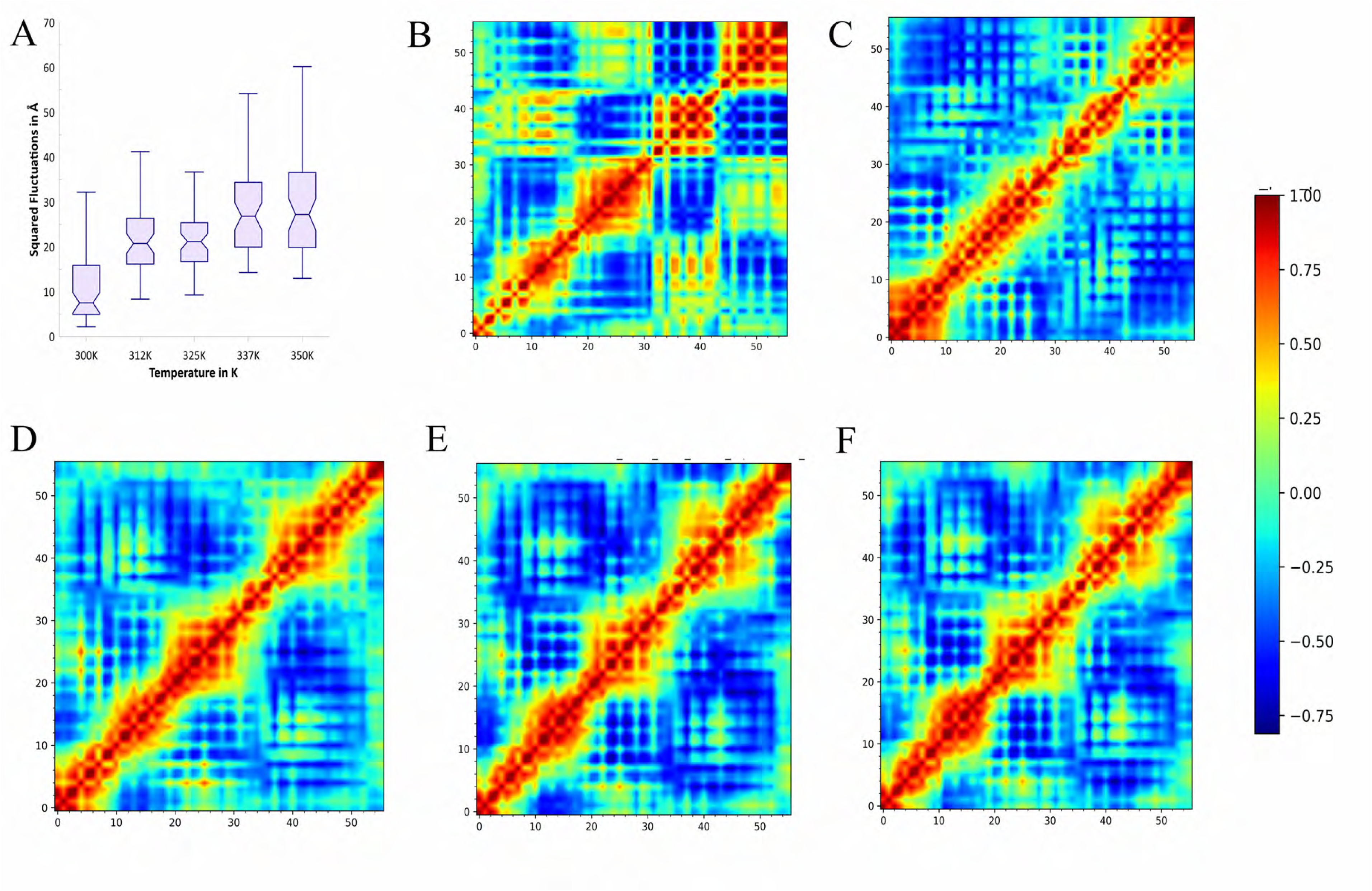
PCA analysis of plot squared fluctuations and cross relations plot for native 2kzd. A) Squared fluctuations in RMSD values at individual temperature profiles as a box plot with pinch at average value. B) to F) Cross relations plot of residues of x-axis and y a-vis for the temperature 300K, 312K, 325K, 337K and 350K respectively. The colour set along with values are provided for all plots.

**Table 1:**
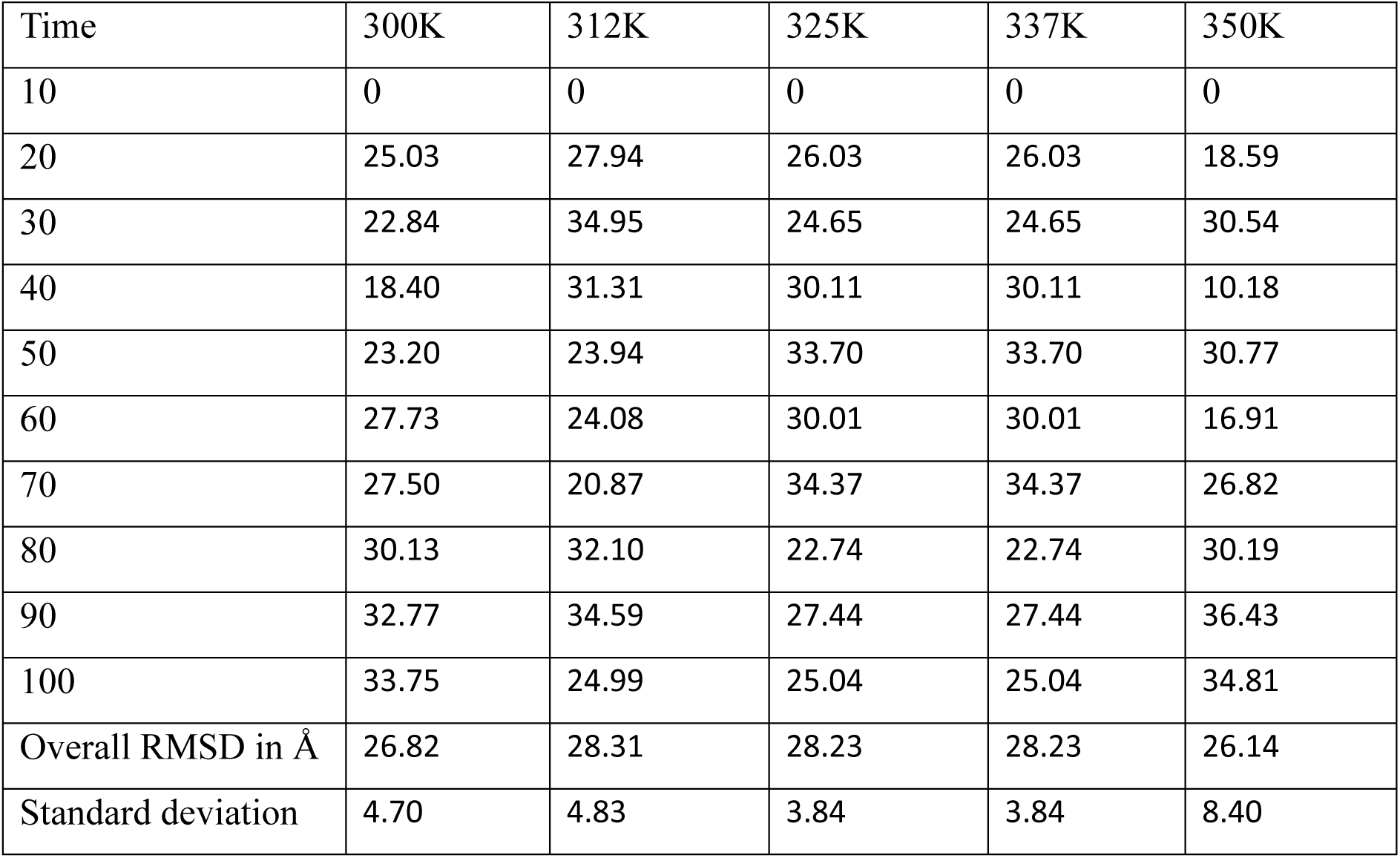
RMSD values of native 2kzd across various temperature profiles.

A similar study was performed for the native structure 2kze to gain insights on the confirmational changes of the quadruplex structure in its native form under the influence of various temperature profiles.

An average RMSD of ∼27 Å, is found to be across the temperature profiles. A PCA analysis for performed to reduce the structural dimensionality and gain insights on the fluctuations.

In the Fig 5A, as in the previous case with rise in temperature the structural deviations increase and the same in provided in table 2. A cross correlation plot at each respective temperature profile from 300 to 350K is shown in Fig 4B to 4F respectively. For 312K and 325K the structural profile looks similar and fairly overlapping is nature. The same relationship was seen for the temperature pairs of 337K and 350K. The RMSD values are comparable and no statistically significant differences (P < 0.05) between various temperature profiles.

**Figure 5:**
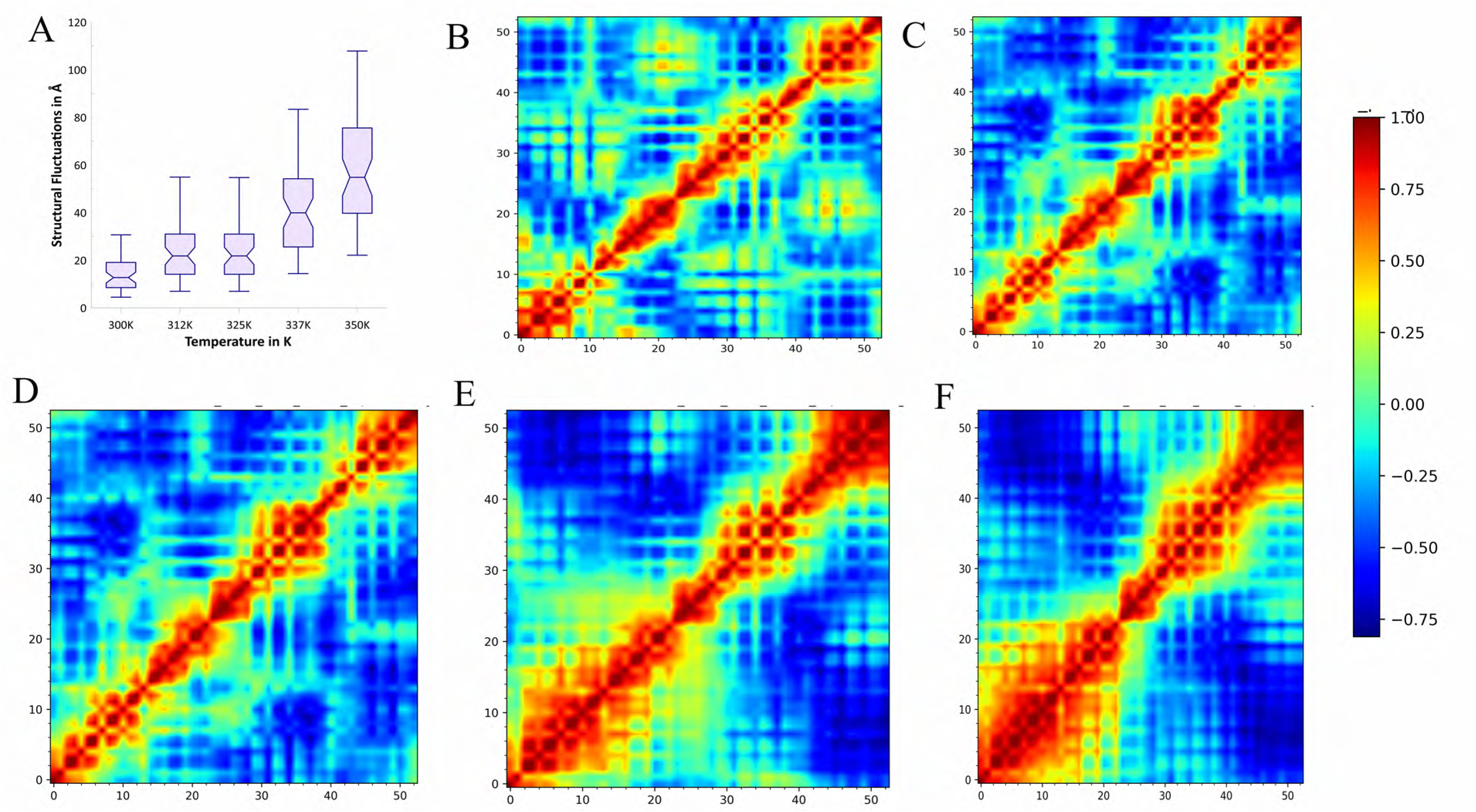
PCA analysis of plot squared fluctuations and cross relations plot for native 2kze. A) Squared fluctuations in RMSD values at individual temperature profiles as a box plot with pinch at average value. B) to F) Cross relations plot of residues of x-axis and y a-vis for the temperature 300K, 312K, 325K, 337K and 350K respectively. The colour set along with values are provided for all plots.

**Table 2:** RMSD values of native 2kze across various temperature profiles.

| Time | 300K | 312K | 325K | 337K | 350K |
| --- | --- | --- | --- | --- | --- |
| 10 | 0.00 | 0.00 | 0.00 | 0.00 | 0.00 |
| 20 | 33.94 | 30.79 | 22.79 | 29.74 | 28.28 |
| 30 | 37.45 | 38.33 | 28.43 | 24.32 | 22.41 |
| 40 | 25.43 | 26.64 | 25.34 | 21.58 | 26.14 |
| 50 | 31.23 | 27.60 | 31.87 | 26.99 | 24.05 |
| 60 | 31.00 | 26.22 | 31.19 | 19.35 | 38.27 |
| 70 | 32.46 | 33.06 | 30.09 | 18.10 | 32.92 |
| 80 | 31.38 | 34.43 | 34.22 | 30.49 | 28.94 |
| 90 | 19.76 | 30.44 | 29.71 | 29.35 | 25.60 |
| 100 | 14.83 | 14.99 | 31.12 | 33.22 | 29.73 |
| Overall RMSD in Å | 28.61 | 29.17 | 29.42 | 25.90 | 28.48 |
| Standard deviation | 6.83 | 6.23 | 3.29 | 5.03 | 4.57 |

#### 3.5.2. REMD studies with control (Maslinic Acid) quadruplex complex

The control compound, Maslinic acid used in the current research with 2kzd was subjected to REMD simulation for 100ns. This will serve as a control study to against the lead compounds.

The RMSD values are in contradiction to the native simulation of the quadruplex structure. The RMSD values in the presence of maslinic acid is ∼ 5Å. The maslinic acid was found to be interacting with quadruplex throughout the simulation period across the various temperature profiles. Even at higher temperature profile the RMSD values is at 5.1Å detailed information provided in table 3. This provides evidence on the stability of the quadruplex structures in the presence of maslinic acid. The RMSD values are comparable and no statistically significant differences (P < 0.05) between various temperature profiles.

**Table 3:** RMSD values of control 1 complex (Maslinic acid with 2kzd) across various temperature profiles.

| Time | 300K | 312K | 325K | 337K | 350K |
| --- | --- | --- | --- | --- | --- |
| 10 | 0.00 | 0.00 | 0.00 | 0.00 | 0.00 |
| 20 | 6.43 | 5.93 | 5.73 | 8.97 | 8.71 |
| 30 | 12.86 | 5.03 | 5.91 | 13.81 | 9.60 |
| 40 | 10.51 | 6.08 | 20.10 | 5.22 | 9.27 |
| 50 | 10.78 | 9.46 | 9.58 | 19.83 | 5.32 |
| 60 | 5.51 | 10.75 | 13.51 | 18.86 | 11.38 |
| 70 | 14.17 | 11.38 | 11.56 | 19.67 | 5.96 |
| 80 | 13.48 | 11.26 | 18.32 | 6.20 | 11.79 |
| 90 | 14.12 | 12.78 | 18.64 | 6.09 | 12.41 |
| 100 | 6.03 | 10.77 | 16.89 | 14.24 | 14.26 |
| Overall RMSD in Å | 6.22 | 6.85 | 4.91 | 5.10 | 5.11 |
| Standard deviation | 1.72 | 1.30 | 0.69 | 1.17 | 1.15 |

The structural profile in fig 6A, it is seen that structural variations are seen to be increasing till 325K and then a decrease in the fluctuation is seen at 337K and 350K. This phenomenon might seem against the conventional understanding of structural conformations under the influence of varied temperature profiles. A discussion on the results seen are justified in the discussion section.

**Figure 6:**
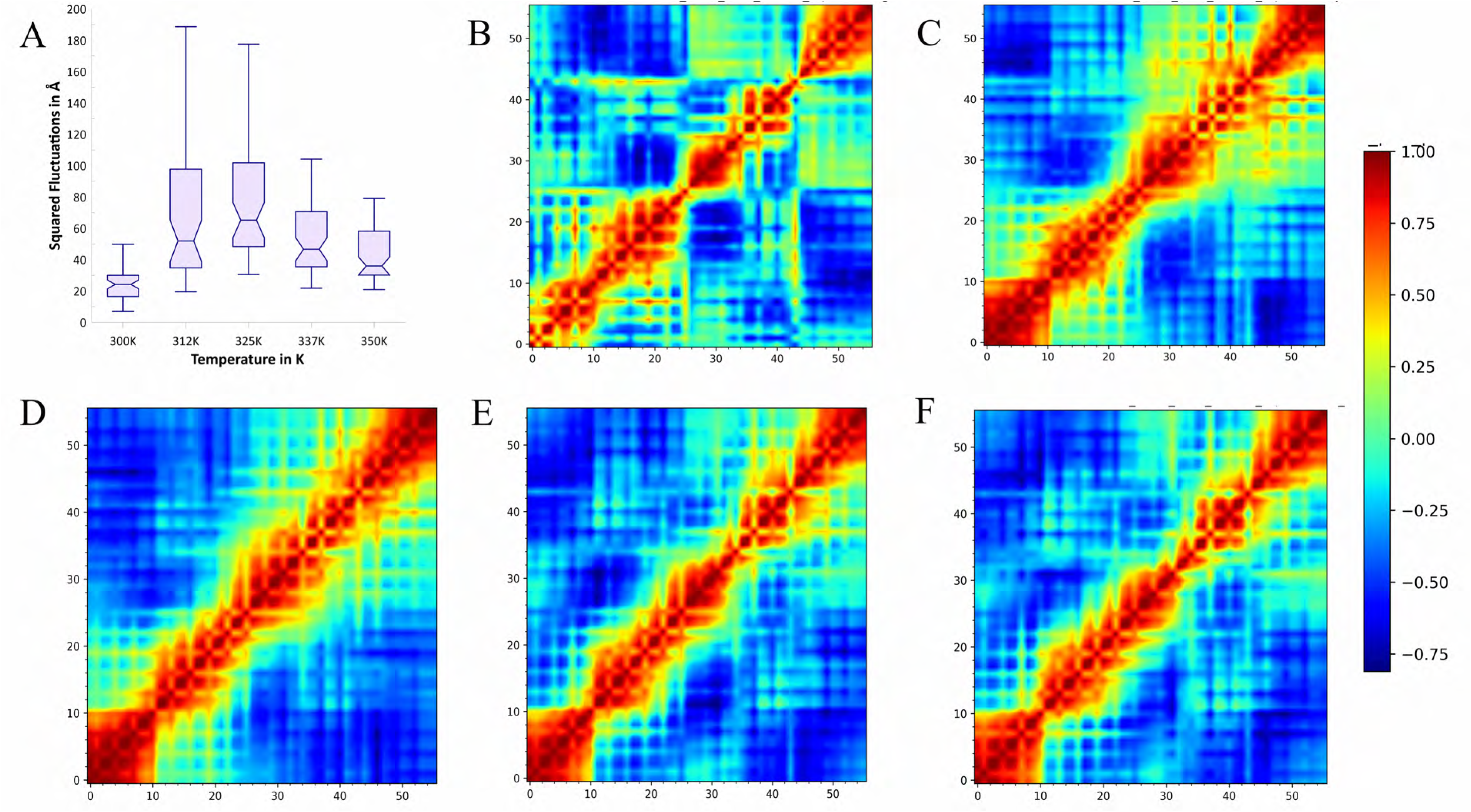
PCA analysis of plot squared fluctuations and cross relations plot for control 1 (Maslinic acid-2kzd). A) Squared fluctuations in RMSD values at individual temperature profiles as a box plot with pinch at average value. B) to F) Cross relations plot of residues of x-axis and y a-vis for the temperature 300K, 312K, 325K, 337K and 350K respectively. The colour set along with values are provided for all plots.

The correlational matrix Fig 6B to 6F for 300K to 350K respectively, shows exactly what has been inferred from the fluctuations where in 312K and 325K show similar structural conformation and 337K and 350K pair have shown less changes in structure conformation and returning to stable 300K structure.

A similar study was performed for the control 2 complex (maslinic acid-2kze) to gain insights on the confirmational changes of the quadruplex structure with maslinic acid at various temperature profiles.

It is clearly seen in the simulation and as well RMSD values tabulated in table 4 that the quadruplex structure is showing behaviour as that of the native 2kze structure. The reason for this replica behaviour is that maslinic acid does not show stable interaction at any temperature profile. The RMSD values are not comparable and there are statistically significant differences (P < 0.05) between various temperature profiles.

**Table 4:** RMSD values of control 2 complex (Maslinic acid with 2kze) across various temperature profiles.

| Time | 300K | 312K | 325K | 337K | 350K |
| --- | --- | --- | --- | --- | --- |
| 10 | 0.00 | 0.00 | 0.00 | 0.00 | 0.00 |
| 20 | 33.94 | 30.79 | 22.79 | 29.74 | 28.28 |
| 30 | 37.45 | 38.33 | 28.43 | 24.32 | 22.41 |
| 40 | 25.43 | 26.64 | 25.34 | 21.58 | 26.14 |
| 50 | 31.23 | 27.60 | 31.87 | 26.99 | 24.05 |
| 60 | 31.00 | 26.22 | 31.19 | 19.35 | 38.27 |
| 70 | 32.46 | 33.06 | 30.09 | 18.10 | 32.92 |
| 80 | 31.38 | 34.43 | 34.22 | 30.49 | 28.94 |
| 90 | 19.76 | 30.44 | 29.71 | 29.35 | 25.60 |
| 100 | 14.83 | 14.99 | 31.12 | 33.22 | 29.73 |
| Overall RMSD in Å | 28.61 | 29.17 | 29.42 | 25.90 | 28.48 |
| Standard deviation | 6.83 | 6.23 | 3.29 | 5.03 | 4.57 |

Maslinic acid get unbound from the structure and shows free movement in the simulation box. Fig 7A perfectly shows the increase in fluctuations of the nucleotides with increase in temperature. This also corresponds to the correlation matrix from Fig 7B to 7F for each temperature profile in increasing order respectively.

**Figure 7:**
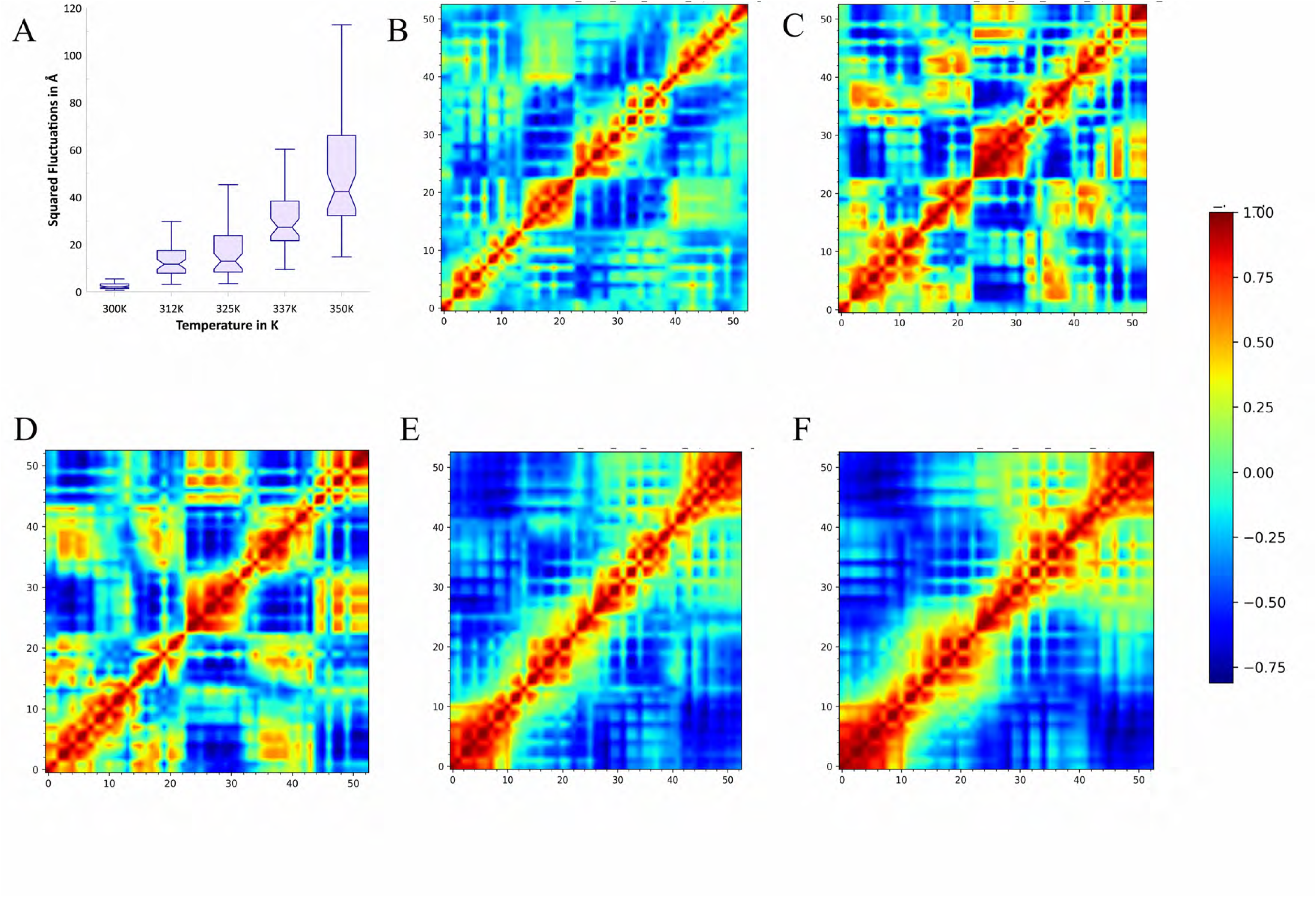
PCA analysis of plot squared fluctuations and cross relations plot for control 2 (Maslinic acid-2kze). A) Squared fluctuations in RMSD values at individual temperature profiles as a box plot with pinch at average value. B) to F) Cross relations plot of residues of x-axis and y a-vis for the temperature 300K, 312K, 325K, 337K and 350K respectively. The colour set along with values are provided for all plots.

This also provides confirmation that maslinic acid does not bind to 2kze conformation of quadruplex and the results are discussed in the next section.

#### 3.5.3. REMD studies with Astragaloside-1 quadruplex complex (2kze)

Astragaloside-1 has shown best docking score with 2kze conformation of quadruplex structure. This complex was subjected to REMD studies as shown in previous sub-sections.

Astragaloside-1 shows extremely high interaction with quadruplex across all the temperature profiles. The fluctuations do not increase beyond 10Å (average of 6Å)in any temperature profile and the deviation value are less. The RMSD values are comparable and no statistically significant differences (P < 0.05) between various temperature profiles. A detailed profile is provided in table 5.

**Table 5:**
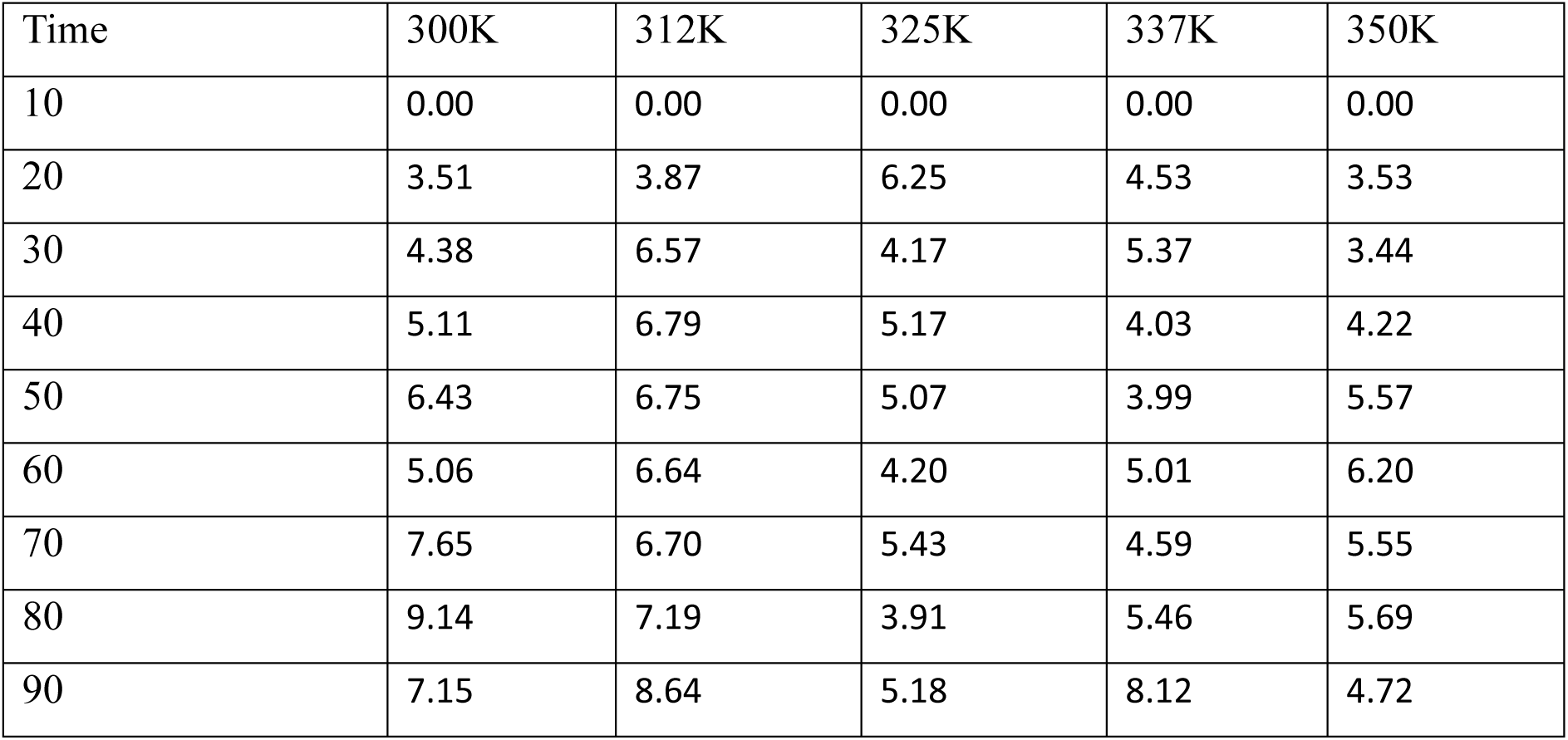
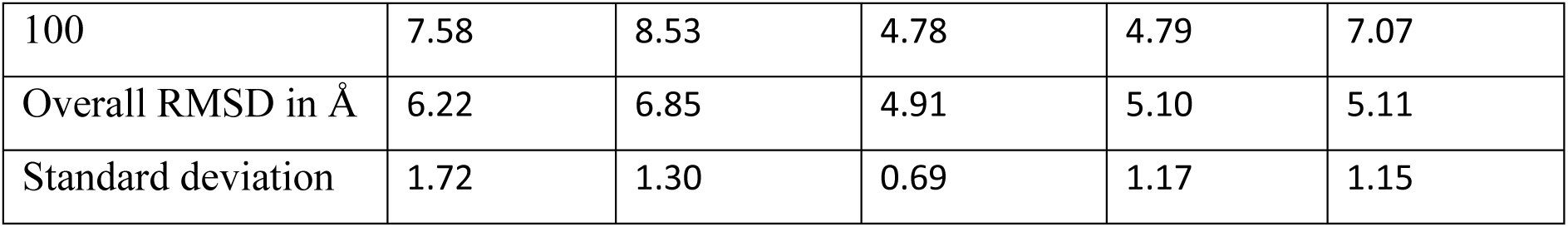
RMSD values of Astragaloside-1 with 2kze across various temperature profiles.

| Time | 300K | 312K | 325K | 337K | 350K |
| --- | --- | --- | --- | --- | --- |
| 10 | 0.00 | 0.00 | 0.00 | 0.00 | 0.00 |
| 20 | 3.51 | 3.87 | 6.25 | 4.53 | 3.53 |
| 30 | 4.38 | 6.57 | 4.17 | 5.37 | 3.44 |
| 40 | 5.11 | 6.79 | 5.17 | 4.03 | 4.22 |
| 50 | 6.43 | 6.75 | 5.07 | 3.99 | 5.57 |
| 60 | 5.06 | 6.64 | 4.20 | 5.01 | 6.20 |
| 70 | 7.65 | 6.70 | 5.43 | 4.59 | 5.55 |
| 80 | 9.14 | 7.19 | 3.91 | 5.46 | 5.69 |
| 90 | 7.15 | 8.64 | 5.18 | 8.12 | 4.72 |
| 100 | 7.58 | 8.53 | 4.78 | 4.79 | 7.07 |
| Overall RMSD in Å | 6.22 | 6.85 | 4.91 | 5.10 | 5.11 |
| Standard deviation | 1.72 | 1.30 | 0.69 | 1.17 | 1.15 |

The squared fluctuation values (Fig 8A) of quadruplex structures with astragaloside-1 shows relative increase (∼12Å) at lower temperatures (300K to 312K) and found to be stable at higher temperatures 325K and beyond. This provides a lot of confidence in the hypothesis of the paper that phytocompounds can assist in stabilization of quadruplex structure across temperatures profiles. The same gets replicated with confidence in the correlation matrix across the temperature profiles from 8C to 8F respectively. A biological interpretation and its implication on the telomerase activity in provided in the discussion section.

**Figure 8:**
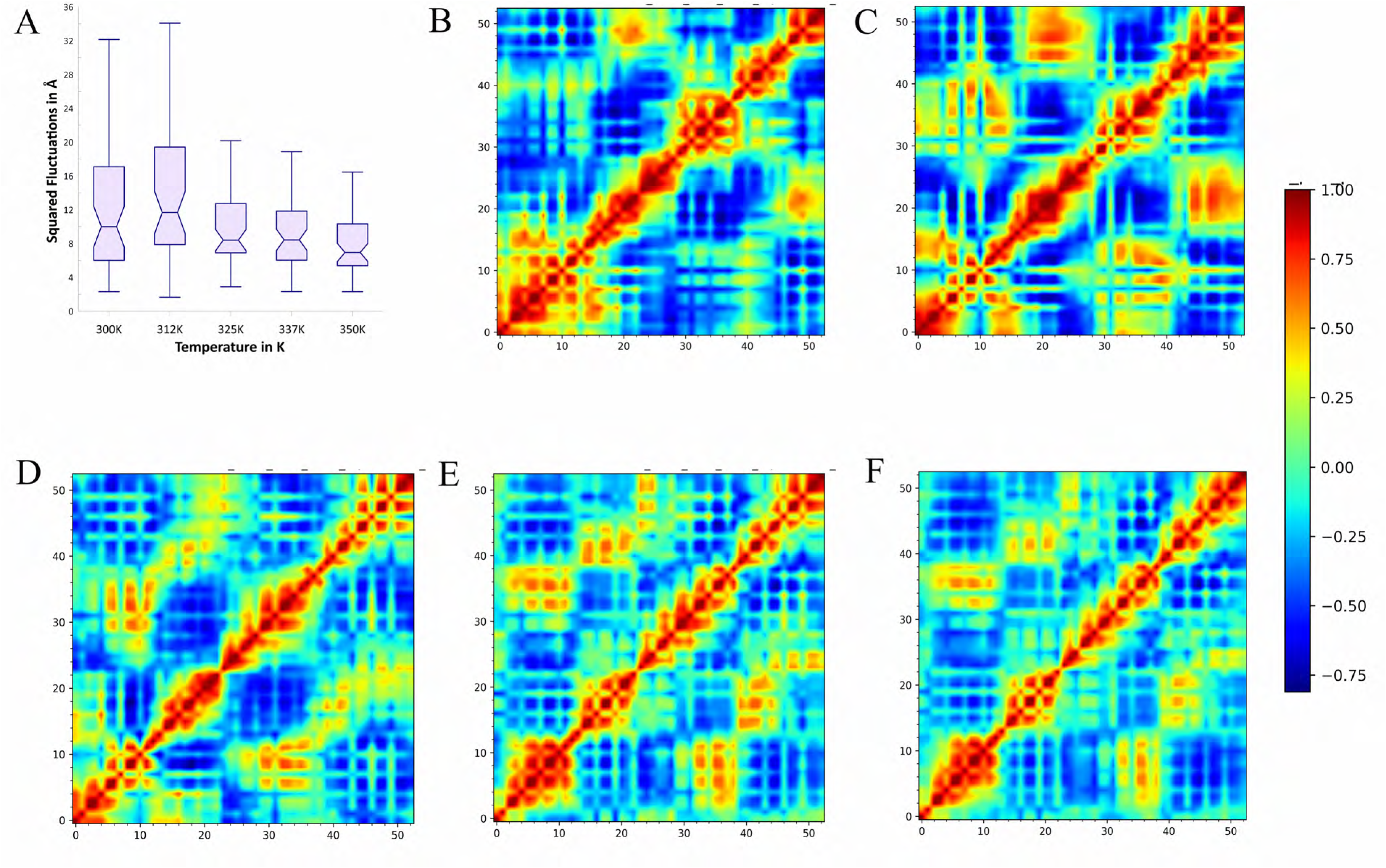
PCA analysis of plot squared fluctuations and cross relations plot for compound 1 (Astragaloside1-2kzd). A) Squared fluctuations in RMSD values at individual temperature profiles as a box plot with pinch at average value. B) to F) Cross relations plot of residues of x-axis and y a-vis for the temperature 300K, 312K, 325K, 337K and 350K respectively. The colour set along with values are provided for all plots.

**Figure 9:**
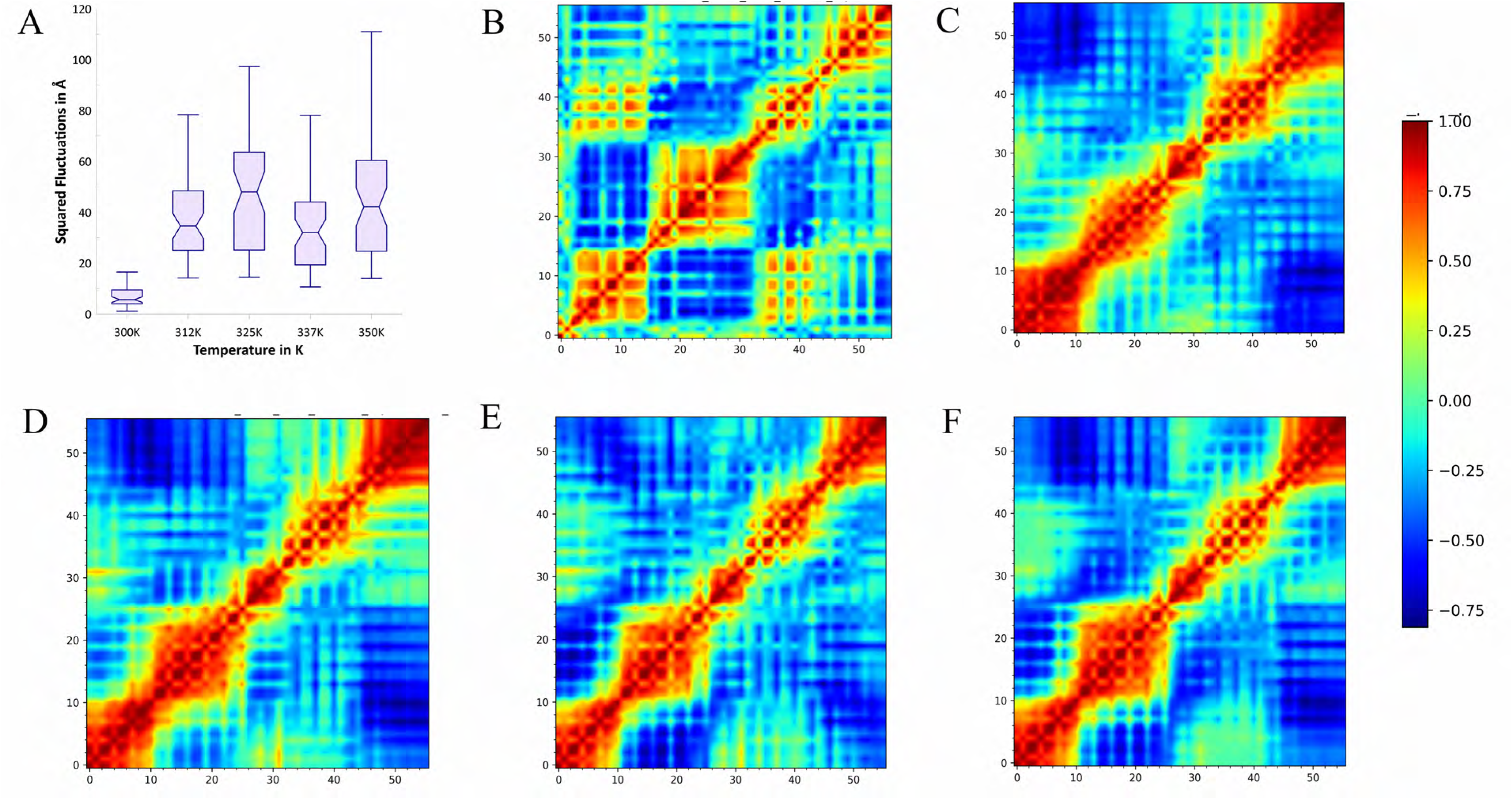
PCA analysis of plot squared fluctuations and cross relations plot for compound 2 (preganone derivative-2kze). A) Squared fluctuations in RMSD values at individual temperature profiles as a box plot with pinch at average value. B) to F) Cross relations plot of residues of x-axis and y a-vis for the temperature 300K, 312K, 325K, 337K and 350K respectively. The colour set along with values are provided for all plots.

#### 3.5.4. REMD studies with pregnane derivative quadruplex complex

Pregnane derivative bearing an IUPAC name (2S)-2-[[4-[2-[(8R,9S,10R,13S,14S,17R)-17-hydroxy-10,13-dimethyl-3-oxo-2,6,7,8,9,11,12,14,15,16-decahydro-1H-cyclopenta[a]phenanthren-17-yl]-2-oxoethoxy]-4-oxobutanoyl]amino]butanedioic acid which showed highest docking score with 2kzd was subjected to REMD simulation.

Pregnane derivative was found to bound tightly with 2kzd at 300K but at other temperature profiles of 312K and beyond it remained unbound and the RMSD values provided in table 6 shows it evidently.

**Table 6:** RMSD values of pregnane derivative with 2kzd across various temperature profiles.

| Time | 300K | 312K | 325K | 337K | 350K |
| --- | --- | --- | --- | --- | --- |
| 10 | 0.00 | 0.00 | 0.00 | 0.00 | 0.00 |
| 20 | 5.91 | 28.67 | 34.67 | 34.56 | 30.75 |
| 30 | 6.77 | 23.12 | 18.51 | 25.49 | 28.77 |
| 40 | 10.01 | 21.61 | 20.44 | 36.02 | 19.71 |
| 50 | 7.09 | 26.97 | 26.32 | 28.51 | 27.66 |
| 60 | 7.25 | 37.18 | 30.69 | 27.02 | 11.70 |
| 70 | 6.74 | 26.14 | 30.18 | 20.00 | 27.06 |
| 80 | 11.38 | 17.82 | 32.07 | 34.58 | 31.01 |
| 90 | 7.93 | 29.29 | 33.59 | 34.92 | 28.39 |
| 100 | 18.15 | 24.95 | 35.50 | 27.79 | 28.95 |
| Overall RMSD in Å | 9.03 | 26.20 | 29.11 | 29.88 | 26.00 |
| Standard deviation | 3.62 | 5.15 | 5.77 | 5.15 | 5.94 |

The simulation and the RMSD values listed in Table 6 make it evident that the quadruplex structure is behaving in a manner similar to the native 2kze structure and with maslinic acid and 2kze. There are statistically significant discrepancies (P < 0.05) across different temperature profiles, and the RMSD values are not comparable.

Pregnane derivative breaks free from its binding groove and moves freely within the simulation box same as in the case of 2kzd. The rise in nucleotide variations with temperature is clearly displayed in Fig. 7A. At 300K, the fluctuations are found to be very stable and increase drastically beyond 312K. This matches the correlation matrix for each temperature profile in ascending order, respectively, from Figs. 7B to 7F.

This further confirms that pregnane derivative does not bind to the quadruplex’s 2kze conformation.

## 4. Discussion

In the previous reported studies [13], the coexistence of two distinct G-quadruplex conformations, Form 1 and Form 2, in the hTERT promoter. The study shows that the G-rich sequence in the hTERT promoter has a high potential to form G-quadruplex structures, and that eight out of eleven candidate sequences containing four consecutive G-tracts can form stable G-quadruplexes. The two G-quadruplex conformations, Form 1 (2kzd) and Form 2 (2kze), were found to be isoenergetic and interconvertible under different experimental conditions, including temperature, pH, and molecular crowding. The study suggests that the coexistence of two distinct G-quadruplex conformations in the hTERT promoter may have functional implications for cancer cell biology and telomere elongation. Form 1 (2kzd) is an intramolecular (3 + 1) G-quadruplex conformation that has been identified in the hTERT promoter sequence. It is favoured at lower temperatures and under certain sequence contexts. The study suggests that in a longer natural sequence context, the formation of Form 1 is possible, indicating its potential stability and relevance within the hTERT promoter. The presence of Form 1 in the hTERT promoter sequence suggests its potential role in telomere biology and cancer cell biology. G-quadruplex structures, including Form 1, have been associated with regulating telomerase activity, affecting gene expression, and influencing the splicing pattern of hTERT RNA. Therefore, the formation of Form 1 may have implications for telomere elongation and cancer cell proliferation.

Form 2 (2kze) is a parallel-stranded G-quadruplex conformation identified in the hTERT promoter sequence. The imino proton patterns suggest the formation of a three-G-tetrad G-quadruplex with a different scaffold, distinguishing it from Form 1. Form 2 is favoured at higher temperatures and under molecular crowding conditions. This indicates its potential stability under specific environmental conditions and suggests its relevance within the hTERT promoter sequence.

In the current study, astragaloside-1 showed stable binding with 2kze and since it is favoured at higher temperatures and the compound stabilized structure which is less than 10Å. This provides a greater confidence in the selection of compound as well as its ability to stabilise the structure even at over molecular crowding scenario.

In continuation to the results section, 2kzd was found to be best docked with preganone derivative but could not stay bound with stable interactions. Even maslinic acid, did not stay stable in REMD simulations. 2kzd adopts an intramolecular G-quadruplex structure with a (3 + 1) topology, characterized by the stacking of three G-tetrads and one edgewise loop. This conformation is stabilized by the Hoogsteen hydrogen bonding [52] within the G-tetrads and the interactions between the loops and the G-tetrads. The major groove of 2kzd in the G-quadruplex structure is involved in specific interactions and recognition processes. The unique arrangement of guanine residues and loop orientations within 2kzd contributes to the distinct architecture of the major groove, which may play a role in ligand binding, protein recognition, and potential functional interactions within the hTERT promoter sequence. This requires specific design of ligands to bind and stabilize the structure.

Overall, in this paper we were able to shortlist Astragaloside-1 as one of the lead phytocompound against the 2kze conformation of hTERT quadruplex.

## 5. Conclusion

The G-quadruplex (G4) structure in the hTERT (human telomerase reverse transcriptase) promoter has gained significant attention in drug discovery. The hTERT promoter’s G-rich region exhibits a high potential for G4 formation, with evidence of the coexistence of distinct G4 conformations. Stabilizing the higher-order hTERT G-quadruplex with small molecules has been shown to be a viable approach.

Astrgaloside-1 has been shown via molecular docking and REMD simulations to exert stable binding and interaction across temperature profiles with 2kze.The same result could not be mimicked for 2kzd as the distinct architecture demands design of novel compounds to stabilize the same.

In the paper, we provide both positive and negative results of compounds which were lead for respective quadruplex structures. The results also show that Insilico simulation techniques are efficient in screening and providing lead compounds against DNA targets accurately.

## 6. Declaration

### 6.1. Author Contributions

V.N. was in for ideation and conceptualization. A.U. contributed to the computational analysis and drafting the manuscript. Both the authors have read and agreed to the published version of the manuscript.

### 6.2. Funding

No funding was available for this work.

## 6.3. Acknowledgement

The authors thank Department of Computer Science and Engineering, the R V College of Engineering, and Bangalore for providing GPU (NVIDIA A100) computational support. A warm heartfelt thanks to the staff and administration at the R V College of Engineering for their support.

## 6.4. Conflict of Interest

The authors declare no conflict of interest.

## Notes

### Competing Interest Statement

The authors have declared no competing interest.

